# Rapid tissue-specific screening of *Wolbachia*, *Cardinium* and *Rickettsia* in Flies (Diptera: Sepsidae; Drosophilidae)

**DOI:** 10.1101/2021.03.11.434924

**Authors:** Cher Wei Yuan, Ding Huicong, Nalini Puniamoorthy

## Abstract

Maternally transmitted endosymbionts can negatively influence the reproduction of their arthropod hosts (e.g. male-killing, cytoplasmic incompatibility). However, such infections are rarely assessed in insect models such as Sepsidae or Drosophilidae that are routinely used in sexual selection studies. To detect infection and bacterial localisation in the host, we developed and optimised a tissue-specific multiplex screening protocol for *Wolbachia*, *Cardinium* and *Rickettsia* that can be completed in a day. The robustness of the protocol was tested with the screening of multiple species and populations of flies commonly used in reproductive studies (N=147 flies; n=426 tissues). With triplex PCRs and more effective duplex PCRs, we detected both single and co-infections in most individuals from both families (Drosophilidae | Sepsidae; Single infection: 51.4% | 62.7%; Dual infection: 29.2% | 9.3%; Triple infection: 4.2% | 0%). Surprisingly, we documented the presence of all three reproductive bacteria in 32 wild-caught drosophilids from Singapore. Also, we note that most sepsid populations (19 out of 22) tested positive for *Cardinium*. We found that the *Rickettsia* infection was overall low, but it was predominantly detected in the gastrointestinal tract instead of the reproductive tract, suggesting a potential horizontal transmission. Finally, we found that amplicon sequences of equivalent sizes between the three tissues from the same individuals share at least 98.8% identity, which suggests that the same endosymbiont strain inhabits within the whole arthropod. Overall, we believe this protocol is effective in detecting co-infections and understanding the transmission of various reproductive endosymbionts. It can also be used to assess endosymbiont infections in other insects.

## Introduction

Reproductive endosymbionts such as *Wolbachia*, *Cardinium* and *Rickettsia* are maternally inherited intracellular bacteria in arthropods (1). Since the production of males are ‘dead ends’ for maternally inherited bacteria, these bacteria often manipulate their host reproduction to favour female offspring (2). For instance, they can induce four possible phenotypes in their host: cytoplasmic incompatibility (CI) between infected sperm and uninfected egg, parthenogenesis in which females emerge from unfertilized eggs, the feminization of genetic males, and the killing of male progeny (2, 3). In addition to these phenotypes, the endosymbionts may also induce host-specific behavioural and immune responses which can influence fitness (3). Furthermore, the phenotype induced in their host varies across bacterial lineages. For instance, all four phenotypes have been associated with *Wolbachia* but the last phenotype, also known as male-killing, is rare (3). In contrast, *Rickettsia* infections are often associated with male-killing (4) and parthenogenesis (5). All but male-killing supposedly occurs as a consequence of *Cardinium* infections (6). Some reports suggest that co-infection by multiple bacteria can influence the severity of CI or even introduce a phenotype that is absent in single infections (7, 8), such as feminization (9). However, relatively little is known about the consequences of co-infections by different bacteria because most studies only screen for one type of bacterium at a time, mostly *Wolbachia*.

The occurrence of reproductively transmitted bacteria likely has consequences for sexual selection and speciation research in arthropods. For instance, studies that investigate barriers to gene flow between two populations may interpret reduced offspring survival of hybrid crosses as an indication of postzygotic reproductive barriers (10). However, CI, which kills a portion of the progeny, may arise in crosses between populations with stark differences in bacterial infection status. As such, CI may confound the results in such studies (10, 11). Even studies on precopulatory courtship and mating behaviour can be affected by the presence of reproductive endosymbionts because these infections may affect mating behaviour and mate choice (12–16). Therefore, researchers investigating sexual selection or speciation in arthropods can benefit by testing the infection status in their study organisms: in-laboratory cultures or strains (where contaminations and infections may spread easily) as well as in wild-caught samples (where the wild populations may be heterogeneously infected) (11,17,18). Here, we screened multiple reproductive bacteria in two families of flies that have been commonly used as model organisms in sexual selection research: Sepsidae and Drosophilidae.

Sepsid flies, also known as black scavenger flies, are dung flies whose larvae play an important role in breaking down organic waste (19). Over the last thirty years, several comparative studies on sepsids suggest that reproductive behaviour and morphology evolves rapidly across species (20–22). In particular, *Sepsis punctum* is an emerging model organism for research on population diversification and incipient speciation (23–25). However, to date, there is no published research investigating possible sepsid infections by reproductive endosymbionts. Drosophilidae represents another family of flies that have been widely used in sexual selection research: from studies on courtship behaviour and cuticular hydrocarbons (12,26–28) to polyandry and sperm competition (29–31). Notably, *Drosophila* spp. were the subject in several landmark papers on reproductive evolution (32–34). There are numerous studies on the presence and consequences of *Wolbachia* in drosophilid flies. For instance, early work investigated the effects of endosymbiont presence and bidirectional CI on divergent evolution of *Drosophila simulans* (35), and recent studies even use *Wolbachia* infections as a biocontrol in regulating *Drosophila suzukii* populations. However, there is little information on the co-infections with other bacteria such as *Rickettsia* and *Cardinium* across multiple species and wild-caught flies.

There is also incomplete information on the mode of transmission of these reproductive bacteria. For instance, although maternal transmission is dominant (3), several studies suggest that endosymbionts might transmit horizontally between individuals and even species (36–40). Much of the current literature on endosymbiont screenings is based on a conventional PCR protocol where the entire arthropod body is macerated before DNA extraction and PCR amplification (10,11,41–44). The mode of transmission, whether vertical, horizontal, or both, cannot be discerned with this method. A solution is to dissect and isolate the somatic and germline tissue of the arthropod which allows inference on the mode of transmission: if the endosymbiont is only detected in the germline, it is likely transmitted vertically down the maternal lineage (39, 45); if found in the gastrointestinal tract, the bacteria may also be transmitted horizontally (39, 46). Additionally, several studies report the integration of bacterial genes into the host genome (47–49). The screening of non-gastrointestinal somatic tissues, such as the muscle, thoracic ganglion, brain or fat body (50), can help detect such cases. This tissue-specific methodology is rarely adopted in published studies because the screening process is protracted by dissection procedures (51). Alternatively, many studies use fluorescence in situ hybridization microscopy to detect localisation (50, 52) but this is often time-consuming and becomes tedious when screening hundreds of specimens.

Hence, we present a new tissue-specific PCR screening protocol to test for the presence of three different reproductive endosymbionts: *Wolbachia*, *Cardinium* and *Rickettsia*. We have used it to test for the presence of multiple infections in the legs (somatic), the gastrointestinal tracts (somatic tissue facilitating horizontal transmission) as well as the reproductive tissues of sepsid and drosophilid flies. Furthermore, we sequenced every amplicon from five individuals with all three tissues infected with either *Wolbachia* or *Cardinium* to examine the possibility of tissue compartmentalization of different strains of the endosymbiont. We believe this study is also relevant to non-fly researchers because this protocol can be optimised to screen other insect models.

## Material and Method

### Species and populations used for endosymbiont screen

For Sepsidae (n=75), we tested individuals from 22 populations of *Sepsis punctum* as well as a congeneric species, *Sepsis orthocnemis*, and two *Dicranosepsis* spp. (sister species to Sepsis) (see Table 1) (22). All cultures were established in the Insectary at National University of Singapore based on previously published protocols (24, 53). For Drosophiliae (n=72): we tested three populations of *Drosophila melanogaster* as well as 12 congeneric species (Table 1). Samples were obtained from the Bloomington *Drosophila* stock centre (Bloomington, IN) as well as the Center for Reproductive Evolution (Syracuse, NY). We also included 32 wild-caught *Drosophila* samples from various locations in Singapore (Fig S1). A total of 147 flies were dissected and 426 tissues were screened.

**Table 1.** Summary of *Wolbachia, Cardinium* and *Rickettsia* infection rate per tissue. Stock flies were collected from the following locations: Bloomington Drosophila stock center (BMT), the Center for Reproductive Evolution (CRE), and the current study’s laboratory culture (LC).

| Species | Country | Collection site | *Wolbachia* infection rate |  |  | *Cardinium* infection rate |  |  | *Rickettsia* infection rate |  |  | N |
| --- | --- | --- | --- | --- | --- | --- | --- | --- | --- | --- | --- | --- |
|  |  |  | (%) |  |  | (%) |  |  | (%) |  |  |  |
|  |  |  | Leg | Ovary/<br>egg | Gut | Leg | Ovary/<br>egg | Gut | Leg | Ovary/<br>egg | Gut |  |
| Sepsid (Laboratory stock culture) |  |  |  |  |  |  |  |  |  |  |  |  |
| *Dicranosepsis*<br>sp. | China | - | 0.0 | 0.0 | 0.0 | 33.3 | 66.7 | 100.0 | 0.0 | 0.0 | 0.0 | 3 |
| *Sepsis*<br>*orthocnemis* | Switzerland | Sörenberg | 0.0 | 0.0 | 0.0 | 0.0 | 33.3 | 33.3 | 0.0 | 0.0 | 0.0 | 3 |
| *Sepsis*<br>*punctum* | Algeria | High altitude | 0.0 | 0.0 | 0.0 | 16.7 | 33.3 | 33.3 | 0.0 | 0.0 | 0.0 | 6 |
| *Sepsis*<br>*punctum* | Algeria | Low altitude | 0.0 | 0.0 | 0.0 | 33.3 | 100.0 | 33.3 | 0.0 | 0.0 | 0.0 | 3 |
| *Sepsis*<br>*punctum* | Canada | Ottawa | 0.0 | 0.0 | 0.0 | 33.3 | 50.0 | 50.0 | 0.0 | 0.0 | 0.0 | 6 |
| *Sepsis*<br>*punctum* | Germany | Ludwigshafen | 0.0 | 0.0 | 0.0 | 66.7 | 33.3 | 33.3 | 0.0 | 0.0 | 0.0 | 3 |
| *Sepsis*<br>*punctum* | Germany | Nordrach | 0.0 | 0.0 | 0.0 | 33.3 | 66.7 | 100.0 | 0.0 | 0.0 | 33.3 | 3 |
| *Sepsis*<br>*punctum* | Italy | Terni | 0.0 | 0.0 | 0.0 | 16.7 | 16.7 | 33.3 | 0.0 | 0.0 | 0.0 | 6 |
| *Sepsis*<br>*punctum* | Scotland | Keltneyburn | 0.0 | 0.0 | 0.0 | 33.3 | 100.0 | 66.7 | 0.0 | 0.0 | 0.0 | 3 |
| *Sepsis*<br>*punctum* | Spain | Tenerife | 0.0 | 0.0 | 0.0 | 0.0 | 0.0 | 0.0 | 0.0 | 0.0 | 0.0 | 3 |
| *Sepsis*<br>*punctum* | Switzerland | Capriasca | 0.0 | 0.0 | 0.0 | 33.3 | 66.7 | 66.7 | 0.0 | 0.0 | 0.0 | 3 |
| *Sepsis*<br>*punctum* | Switzerland | Zürich | 0.0 | 0.0 | 0.0 | 66.7 | 66.7 | 100.0 | 0.0 | 0.0 | 66.7 | 3 |
| *Sepsis*<br>*punctum* | U.S.A. | Vermont | 0.0 | 0.0 | 0.0 | 33.3 | 66.7 | 100.0 | 0.0 | 0.0 | 0.0 | 3 |
| *Sepsis*<br>*punctum* | U.S.A. | Davis | 0.0 | 0.0 | 0.0 | 0.0 | 0.0 | 33.3 | 0.0 | 0.0 | 0.0 | 3 |
| *Sepsis*<br>*punctum* | U.S.A. | Fargo | 0.0 | 0.0 | 0.0 | 33.3 | 66.7 | 33.3 | 0.0 | 0.0 | 0.0 | 3 |
| *Sepsis*<br>*punctum* | U.S.A. | Georgia | 0.0 | 0.0 | 0.0 | 33.3 | 33.3 | 66.7 | 0.0 | 0.0 | 0.0 | 3 |
| *Sepsis*<br>*punctum* | U.S.A. | Park City | 0.0 | 0.0 | 0.0 | 100.0 | 66.7 | 100.0 | 0.0 | 33.3 | 33.3 | 3 |
| *Sepsis*<br>*punctum* | U.S.A. | Syracuse | 0.0 | 0.0 | 0.0 | 33.3 | 0.0 | 0.0 | 0.0 | 0.0 | 0.0 | 3 |
| *Sepsis punctum* | U.S.A. | Wolverton | 0.0 | 0.0 | 0.0 | 0.0 | 33.3 | 33.3 | 0.0 | 0.0 | 0.0 | 3 |
| **Sepsid (Wild-caught)** |  |  |  |  |  |  |  |  |  |  |  |  |
| *Dicranosepsis* sp. | Singapore | Pasir Ris | 33.3 | 33.3 | 100.0 | 0.0 | 0.0 | 33.3 | 0.0 | 0.0 | 0.0 | 3 |
| *Dicranosepsis* sp. | Singapore | Tampines Eco-green | 100.0 | 100.0 | 66.7 | 0.0 | 0.0 | 33.3 | 0.0 | 0.0 | 0.0 | 3 |
| *Dicranosepsis* sp. | Singapore | Gangsa trail | 0.0 | 0.0 | 0.0 | 33.3 | 33.3 | 100.0 | 0.0 | 0.0 | 33.3 | 3 |
| **Drosophilid (Laboratory stock culture)** |  |  |  |  |  |  |  |  |  |  |  |  |
| *Drosophila bifurca* | - | CRE | 0.0 | 0.0 | 0.0 | 33.3 | 0.0 | 0.0 | 0.0 | 0.0 | 0.0 | 3 |
| *Drosophila hydei* | - | CRE | 0.0 | 0.0 | 0.0 | 100.0 | 66.7 | 33.3 | 0.0 | 0.0 | 0.0 | 3 |
| *Drosophila kikkawai* | - | BMT | 0.0 | 0.0 | 0.0 | 66.7 | 100.0 | 33.3 | 0.0 | 0.0 | 0.0 | 3 |
| *Drosophila melanogaster* | U.S.A. | Carolina, LC | 100.0 | 100.0 | 100.0 | 33.3 | 33.3 | 33.3 | 0.0 | 0.0 | 0.0 | 3 |
| *Drosophila melanogaster* | - | CRE | 100.0 | 100.0 | 100.0 | 83.3 | 50.0 | 66.7 | 0.0 | 0.0 | 16.7 | 6 |
| *Drosophila mexicana* | - | BMT | 0.0 | 0.0 | 0.0 | 50.0 | 50.0 | 50.0 | 0.0 | 0.0 | 0.0 | 2 |
| *Drosophila mojavensis* | - | BMT | 0.0 | 0.0 | 0.0 | 33.3 | 0.0 | 0.0 | 0.0 | 0.0 | 0.0 | 3 |
| *Drosophila novamexicana* | - | BMT | 33.3 | 33.3 | 66.7 | 33.3 | 66.7 | 66.7 | 0.0 | 0.0 | 0.0 | 3 |
| *Drosophila obscura* | - | BMT | 0.0 | 0.0 | 0.0 | 50.0 | 0.0 | 0.0 | 0.0 | 0.0 | 0.0 | 2 |
| *Drosophila pseudoobscura* | - | BMT | 0.0 | 0.0 | 0.0 | 33.3 | 33.3 | 33.3 | 0.0 | 0.0 | 33.3 | 3 |
| *Drosophila takahashii* | - | BMT | 33.3 | 0.0 | 0.0 | 33.3 | 66.7 | 66.7 | 0.0 | 0.0 | 0.0 | 3 |
| *Drosophila virilis* | - | CRE | 0.0 | 0.0 | 0.0 | 33.3 | 33.3 | 66.7 | 0.0 | 0.0 | 0.0 | 3 |
| *Drosophila yakuba* | - | CRE | 33.3 | 66.7 | 33.3 | 66.7 | 66.7 | 66.7 | 33.3 | 33.3 | 33.3 | 3 |
| **Drosophilid (Wild-caught)** |  |  |  |  |  |  |  |  |  |  |  |  |
| *Drosophila ananassae* | Singapore | Hougang | 100.0 | 100.0 | 100.0 | 100.0 | 100.0 | 100.0 | 0.0 | 0.0 | 0.0 | 3 |
| *Drosophila ananassae* | Singapore | Jurong West | 100.0 | 100.0 | 100.0 | 50.0 | 50.0 | 0.0 | 0.0 | 0.0 | 0.0 | 2 |
| *Drosophila ananassae* | Singapore | Nee Soon | 100.0 | 100.0 | 100.0 | 100.0 | 80.0 | 80.0 | 0.0 | 0.0 | 0.0 | 5 |
| *Drosophila ananassae* | Singapore | River Valley | 100.0 | 66.7 | 100.0 | 0.0 | 0.0 | 0.0 | 0.0 | 0.0 | 0.0 | 3 |
| *Drosophila ananassae* | Singapore | Serangoon | 40.0 | 80.0 | 100.0 | 60.0 | 20.0 | 20.0 | 0.0 | 0.0 | 0.0 | 5 |
| *Drosophila ananassae* | Singapore | Tampines | 0.0 | 25.0 | 0.0 | 0.0 | 0.0 | 0.0 | 0.0 | 0.0 | 0.0 | 4 |
| *Drosophila ananassae* | Singapore | Woodlands | 66.7 | 66.7 | 66.7 | 66.7 | 33.3 | 66.7 | 33.3 | 0.0 | 0.0 | 3 |
| *Drosophila melanogaster* | Singapore | Tampines | 0.0 | 0.0 | 0.0 | 0.0 | 0.0 | 0.0 | 0.0 | 0.0 | 0.0 | 1 |
| *Drosophila* sp. | Singapore | Choa Chu Kang | 33.3 | 33.3 | 0.0 | 0.0 | 0.0 | 0.0 | 0.0 | 0.0 | 0.0 | 3 |
| *Drosophila* sp. | Singapore | Jurong West | 100.0 | 0.0 | 100.0 | 0.0 | 0.0 | 0.0 | 0.0 | 0.0 | 0.0 | 1 |
| *Drosophila* sp. | Singapore | Serangoon | 100.0 | 100.0 | 100.0 | 100.0 | 100.0 | 100.0 | 0.0 | 0.0 | 0.0 | 1 |
| *Drosophila* sp. | Singapore | Tampines | 0.0 | 100.0 | 0.0 | 0.0 | 0.0 | 0.0 | 0.0 | 0.0 | 0.0 | 1 |

### Wild-caught *Drosophila*: Sampling and species identification

*Drosophila* spp. were caught in the urban residential areas of Singapore (Table S1 & Fig S1) using a 100 g of yeast-fermented banana as bait (54). We took high-resolution images of the flies (lateral and dorsal) and compared them to digital images of *Drosophila* spp. on “The Biodiversity of Singapore” (55) for putative identification to morphospecies (see http://www.reprolabnus.com/urops-wy.html). We also amplified the *COI* barcoding fragment for molecular species identification (Table S1). *COI* sequences were queried in NCBI BLASTN and BOLD (56).

### Tissue-specific dissection and molecular analysis

We developed a step-by-step protocol to perform tissue-specific dissection and DNA extraction for multiplex PCR amplification. There are a few key points to observe when carrying out the protocol. Firstly, to ensure accurate localization of endosymbionts, the dissected samples must not cross-contaminate one another. In addition to aseptic technique, adherence to a sequence of dissection steps must be observed: first, isolate the legs to prevent it from being contaminated by any bacteria residing within the fly. Next, dissect for the germline tissues without rupturing the gastrointestinal tract (Fig 1). Failure to do so releases gut bacteria into the dissecting solution.

**Fig 1.**
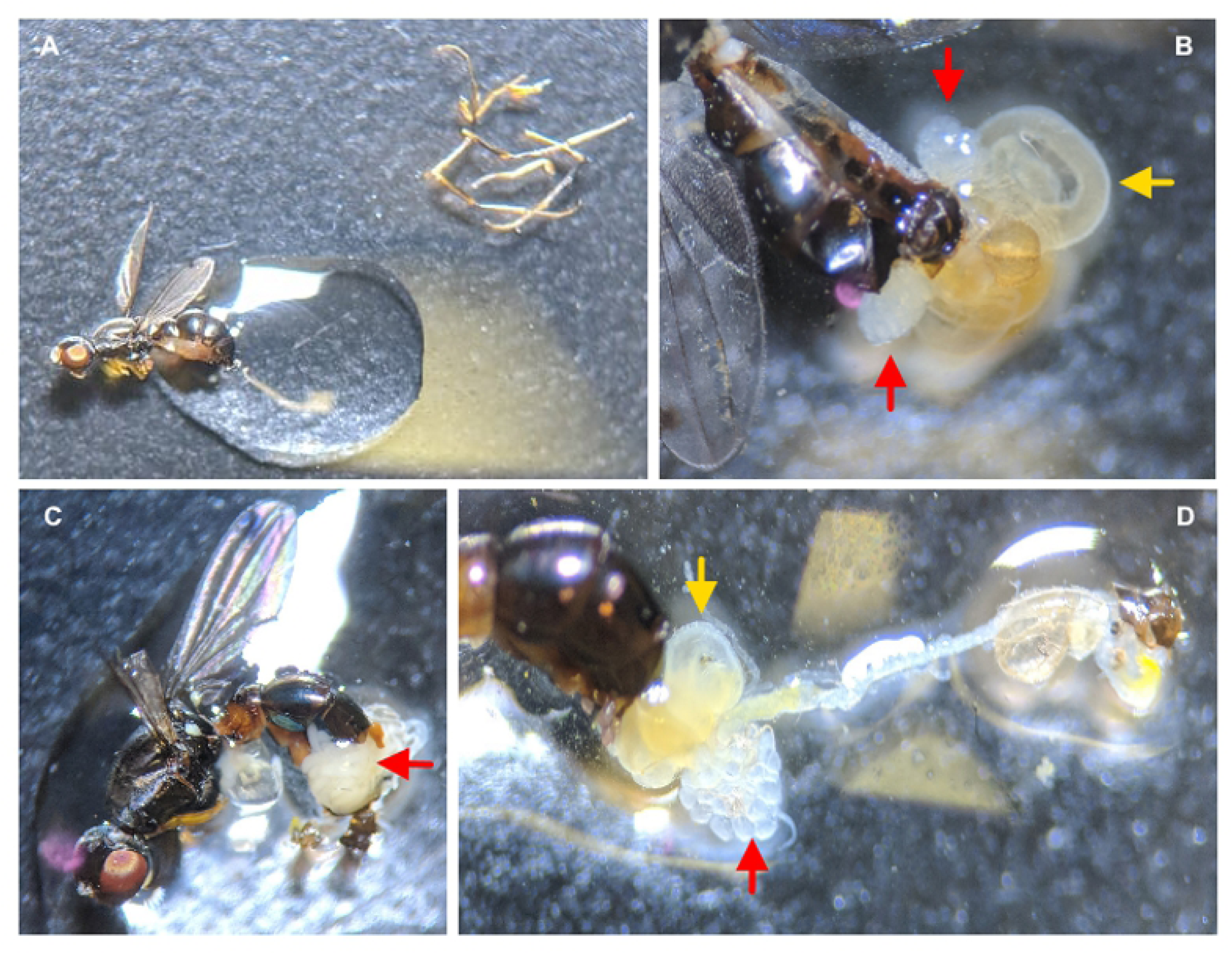
**Tissue-specific dissection of a sepsid fly**. (A) Sepsid fly abdomen placed in water droplet before dissection. Legs are removed before any internal organ dissection to prevent cross-contamination by internal tissues. For reproductive organs (red arrows) dissection: either (B) ovaries or (C-D) eggs were released by slowly slitting the lateral side of the abdomen. After the germline tissue is removed, the gut (yellow arrow) can then be isolated.

Secondly, to expedite the screening, we opted for Lucigen’s QuickExtract^TM^ (QE) to extract DNA because QE allows for rapid extraction of DNA within 30 minutes. The manufacturer recommends using 0.5 mL of QE solution per sample, but we found that a drastically lower volume of 3-5 µL was optimal for the suggested insect tissue sizes (details in protocol): 3 µL for drosophilids and 5 µL for sepsids. Unfortunately, a drawback of QE is the short shelf life of extracted DNA. We found that some QE samples kept for a year could not be successfully amplified by PCR. If long term storage of extracted DNA samples is required, we suggest opting for phenol-chloroform precipitation or kit-based extraction instead.

Lastly, one triplex and two types of duplex PCRs were optimized to screen *Wolbachia*, *Cardinium,* and *Rickettsia* with the primers listed in Table S2. We chose primer combinations with ΔG more than negative one to reduce the risk of primer dimer formation. The calculation of ΔG was done with PrimerPooler (Brown et al., 2017). Subsequently, we optimized a triplex PCR for the *Wolbachia* surface protein gene (*wsp*) as well as the *16S* primers specific for all three bacteria (Table S3, Fig S2) using a mixture of amplified DNA from three positive controls, each infected with *Wolbachia* (*Nasonia* samples from the Werren lab, Rochester, NY), *Cardinium*, and *Rickettsia* (whitefly samples from the Ghanim lab, Volcani Center, Israel). To cross-check the results of the triplex PCR, two duplex PCRs were developed with primers (1) W-spec and *COI* (LCO1498 and HCO2198), where the latter serves as an internal control, and *Car16S* and *Rb16S*. A critical aspect of triplex optimization lies in the balancing of primer concentrations (Sint et al., 2012): we found that 0.2 M of *wsp* and 0.3 M of *Car16S*/*Rb16S* primers were optimal to produce three clear amplicon bands in our positive control (Fig S2). For the duplex reaction, we resolved the differences between the annealing temperature of the primers, W-spec (47 °C) and *COI* (55 °C) for a single PCR via a step-up reaction (47 °C for 15 cycles followed by 55 °C for 20 cycles, see Table S3).

Using the dissection and multiplex protocols, we screened sepsid and drosophilid samples. We cleaned up their PCR products with SureClean Plus (Bioline) and sent 80 amplicons for Sanger sequencing by Axil Scientific (Singapore). We also sent 21 *COI* barcodes for Next-Generation Sequencing to confirm species barcodes (Meier et al., 2015). We trimmed the forward and reverse Sanger sequences with ChromasPro (60) and assembled them with DNA Baser Assembler v3.5.4 (Heracle Biosoft, Arges, Romania). We queried the sequences in NCBI BLASTN.

For phylogenetic analyses, we supplemented our sequences with additional ones from relevant literature (41, 61) and Genbank (62). We performed the analyses on Phylemon web server (63). The sequences were aligned using MUSCLE v. 3.7 (64) with “kmer4_6” and “pctidkimura” as distance measures for iteration one and two respectively. We manually removed sequences on 5’ and 3’ ends that were found only in the supplemental sequences to allow the equal representation of all sequences. Next, we used trimAl (65) on automatic setting to trim the sequences before constructing the trees using MEGAX (66) with the following settings: neighbour-joining tree, 1000 bootstraps, Kimura 2-parameter model, d: transitions + transversions and uniform rates, partial deletion (95%).

### Protocol

#### 1. Before dissection

A. Purify water with MILLI-Q® IQ 7000 and autoclave it at 121 °C for 2 hours.
B. Wear gloves disinfected with 70% ethanol. Disinfect the microscope stage with DNA AWAY^TM^ (Thermo Fisher Scientific) or 70% ethanol.

#### 2. Immobilising flies for dissection

A. Isolate female flies with turgid abdomen from the stock culture into a separate container using a fly aspirator or Pasteur pipette. A turgid abdomen indicates the presence of matured eggs.
B. Freeze the flies in -40°C fridge for 5 to 10 minutes until they are immobilized.
C. Keep the flies on ice.

#### 3. Dissection (5-10 minutes per sample)

A. Sterilize the forceps and dissecting slides with DNA AWAY^TM^ or 70% ethanol. Wipe dry.
B. Place immobilised fly on the cleaned slide.
C. With one forceps holding the thorax, remove all the legs with another forceps (Fig 1A). Set aside the legs in a clean PCR tube. Important note: Cut leg into the femur, tibia, tarsus and macerate them to improve DNA extraction using Lucigen’s QuickExtract^TM^ (QE).
D. Clean forceps with 70% ethanol.
E. Add a small droplet of water with a diameter of 0.5 cm. Place only the abdomen of the insect in the droplet (Fig 1A).
F. Pinch the epithelial on the lateral side of the abdomen with both forceps. Pull the epithelial in opposite directions. The eggs or ovaries should be released from the torn abdomen (Fig 1B-C). Alternatively, squeeze the middle of the abdomen gently to expose the ovipositor using a pair of forceps. With another pair of forceps, grip the ovipositor and pull slowly. In an ideal case, the eggs or ovaries slides out with the gut (Fig 1D). If not, continue pulling until the eggs or ovaries are exposed.
G. Place 2 eggs for sepsid-size insect (3 eggs for Drosophila-size insect) into a new PCR tube. If required, view under the microscope to ensure the eggs has been deposited into the tube. A helpful tip to ease egg/ovary collection: carry the eggs with a small droplet of water in between the tip of the forceps to prevent the tissue from sticking to the forceps. Important note: If the gut is broken in this step, as indicated by the water droplet turning turbid, discard sample or use it for whole body DNA extraction. If long-term storage of the extracted DNA is required, opt for other DNA extraction methods (e.g. column purification).
H. Clean forceps with 70% ethanol. Wipe dry.
I. Cut a segment of the gut the length of the femur of the sepsid (and tibia for Drosophila) and place in new PCR tube. Clean forceps and wipe dry before handling another sample.
J. Place dissected tissues on ice immediately to prevent degradation. Leave the lid open for 5 minutes under a clean cover to remove any residual ethanol. Store in -20°C.

#### 4. DNA extraction

A. Add 5 µL of QE solution for a sepsid-size specimen or 3 µL for Drosophila-sized sample to each PCR tube. Centrifuge for 5 seconds.
B. Place the tubes into the thermocycler to heat at 65°C for 18 minutes.
C. Cool the tubes on ice for 10 minutes and spin down for 5 seconds using a tabletop centrifuge.

#### 5. PCR and agarose gel electrophoresis

A. Duplex PCR: Make a master mix for PCR with each tube containing 12.5 µL of OneTaq® Quick-Load® 2X Master Mix with Standard Buffer, 1 µL of 1 mg mL⁻^1^ BSA, 0.184 µL of 25mM magnesium chloride and 2 µL of each of the two primers diluted to 5 µM (refer to Table S3). Add 1-2 µL of extracted DNA. Top up the final mixture to 25.0 µL. Triplex PCR: In each PCR tube, add 12.5 µL of OneTaq® Quick-Load® 2X Master Mix with Standard Buffer, 1 µL of 1 mg mL⁻^1^ BSA, 0.184 µL of 25mM magnesium chloride, 1 µL of each of the *Wsp* primers and 1.5 µL of each of *Car16S* and *Rb16S* primers (all primers diluted to 5 µM). Add 1-2 µL of extracted DNA. Top up the final mixture to 25.0 µL. Important note: The triplex may be reliable than the duplex PCR in certain species/populations (see results), so, it is recommended to test triplex with positive controls before employing it for large scale screening.
B. Run PCR with cycle conditions specified in Table S3. Remember to include appropriate positive and negative controls.
C. Cast a 2% agarose gel with 1.5 µL GelRed per 40 mL of 1X TAE buffer. Load 1.5-2 µL of sample and control PCR products. Conduct gel electrophoresis for 40 minutes at 90V. Visualize DNA bands using a UV transilluminator.

### Troubleshooting

Refer to Supplementary Table S4.

### Statistical analyses

We conducted Cochran’s Q test to identify any significant difference in the proportion of infection among the three tissues for each of the primers, *W-spec*, *Wsp*, *Car16S* and *Rb16S*. McNemar’s post-hoc analysis was performed as a follow up to identify tissue pairs that differed significantly in infection proportion. Individuals without the internal control (*COI* gene) amplified for all three tissues were excluded from the statistical analysis, leaving n = 71 dissected individuals suitable for analysis. The exclusion of such samples ensures that tissues tested negative for bacteria were likely due to the absence or low count of endosymbionts, and not the failure of PCR. The same dataset was then analysed for differences in the proportion of each bacteria detected between duplex and triplex PCR using Fisher’s Exact test and Chi-square test with Yates’ continuity correction. Statistics were done in RStudio (67) with packages nonpar (68) and rcompanion (69).

To examine whether the same endosymbiont infects different tissues of the same individual, we sequenced *Wolbachia* and *Cardinium* 16S amplicons from leg, egg and gut of five individuals. We aligned these using the Smith-Waterman algorithm (gap open = 10, gap extension = 0.5) in EMBL-EBI. A high similarity would imply the same strain of bacteria across the tissues.

## Results

### Effectiveness of the protocol

The proportion of infection among the three dissected tissues was statistically different for the *Rickettsia* primer (*Rb16S*) (Q = 12, p-value < 0.01). The proportion of infection in the gastrointestinal somatic tissue was significantly different compared to either the germline or appendage tissue (χ^2^ = 6, adjusted p-value < 0.05).

Interestingly, the triplex PCR was unable to detect *Rickettsia* infection in individuals (n = 71) despite amplification of internal control for all three tissues. In contrast, in the same samples, the duplex PCR detected seven infected individuals which was significantly different from the triplex (χ^2^ = 45.8, p-value < 0.001). Similarly, the duplex PCR found 45 individuals infected with *Cardinium* but only ten were detected by the triplex PCR in the same samples. *Cardinium* detection was not statistically different between the two PCRs (p-value = 0.31, Fisher’s Exact test). For *Wolbachia* detection, the duplex fared significantly better (p-value < 0.001, Fisher’s Exact test).

### Infection status of sepsids and drosophilids

Based on 426 tissue-specific screens (Table 1), we found major differences in the infection status of *Wolbachia*, *Rickettsia* and *Cardinium* in both families of flies (No infections: *Drosophila* = 15.3 %, Sepsidae = 28.0 %; Single infection: *Drosophila* = 51.4 %, Sepsidae = 62.7 %; Dual infection: *Drosophila* = 29.2 %, Sepsidae = 9.3 %; Triple infection: Sepsidae = 0.0 % *Drosophila* = 4.2 %; also see Table S5). Overall, *Rickettsia* infection was low (*Drosophila* = 5.6 %, Sepsidae = 6.7 %). Interestingly, all samples with *Rickettsia* infection also had *Cardinium* (Table 1). A subset of amplicons was sequenced and deposited in GenBank under the accession number: MN864660- MN864706 and MN900912-MN900950 (Table S6 & S7). The only sepsid samples providing positive bands of expected size from *Wsp* primers were from Singapore. However, each sequence only has an approximate 30 bp alignment with their respective top three BLASTN results which were *Wolbachia* sequences (Table S8). We translated the five sequences and performed multiple alignments: only 6 out of 188 sites were polymorphic. However, these protein sequences did not match any *Wolbachia* with BLASTP. Taken together, the evidence suggests that Singapore sepsids harboured a different strain of *Wolbachia* or another species of bacteria related to *Wolbachia*. Almost every population (except for Tenerife, Wolverton and Tampines Eco-Green park) exhibited infections of *Cardinium*, and 66.7% of all sepsid individuals gave positive PCR results with *Car16S* primers (Table S5; BLAST results in S8). However, in the drosophilid samples, *Wolbachia* was detected mainly in the wild-caught individuals but the stock cultures were predominantly infected by *Cardinium* (Table S5; BLAST results in S9).

To check if the amplicons of the same size were derived from the same bacterial species, we sequenced amplicons of all three tissues from 6 individuals and aligned pairwise the three sequences from each individual (Table 2). We use a threshold of 98.7% average nucleotide identity (ANI) to classify bacteria as the same species (70). Our results show that the lowest ANI was 98.8% (Table 2) which suggest that each dissected tissue from an individual was infected by the same bacteria species. The amplicon sequences were also queried in BLASTN. Most of the *Cardinium* 16S sequences obtained from sepsid populations had the same top BLAST results: *Cardinium* of whitefly *Bemisia tabaci* (Table S8). However, this could be due to the low number of *Cardinium* sequences in GenBank. For instance, we conducted an extensive search of GenBank with the term ‘*Cardinium*’ (1356 hits) and analysed each result individually. We discovered that only 765 of these sequences were *Cardinium* from screening studies. In contrast, there were 204,229 *Wolbachia* sequences.

**Table 2.**
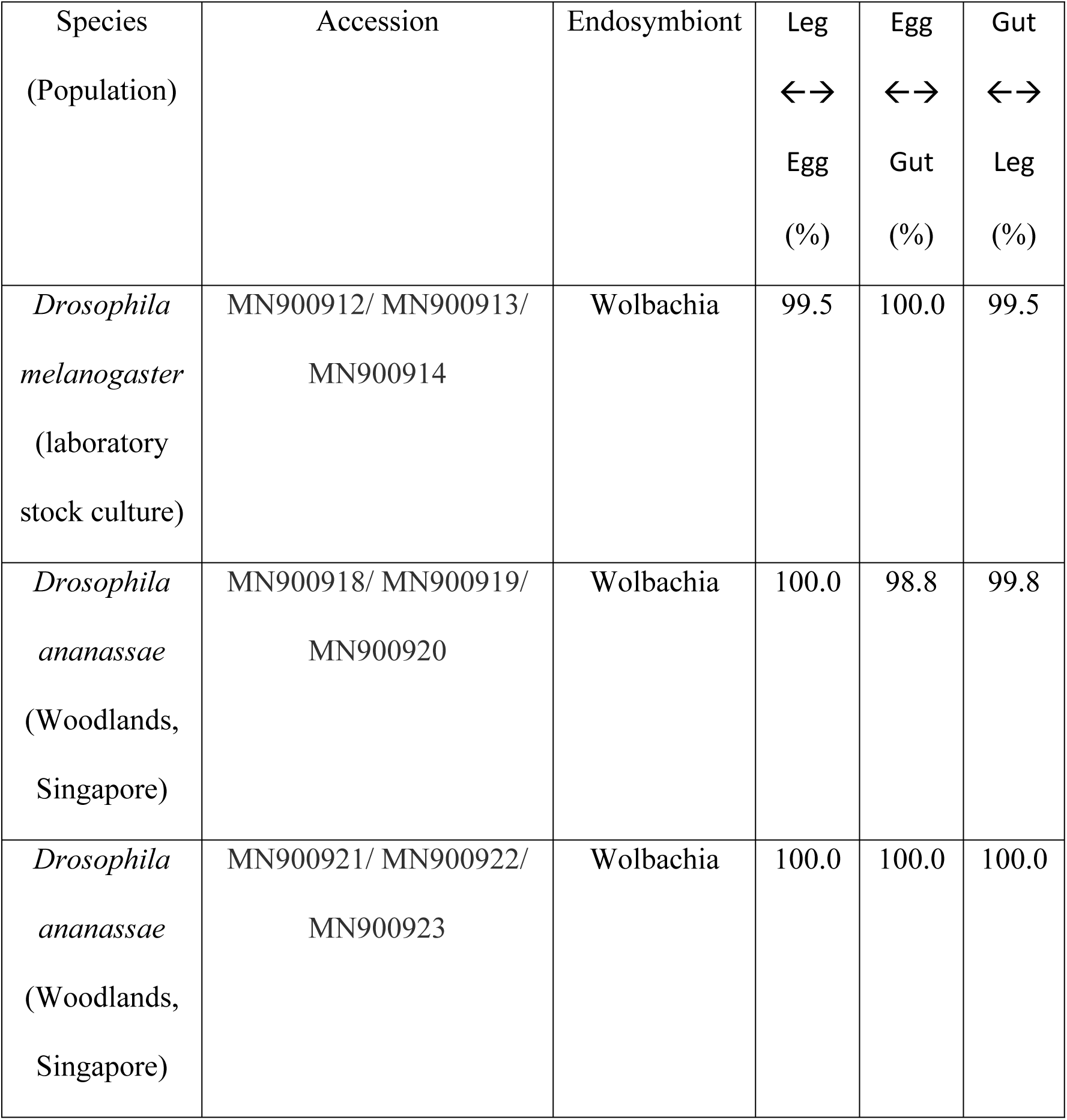
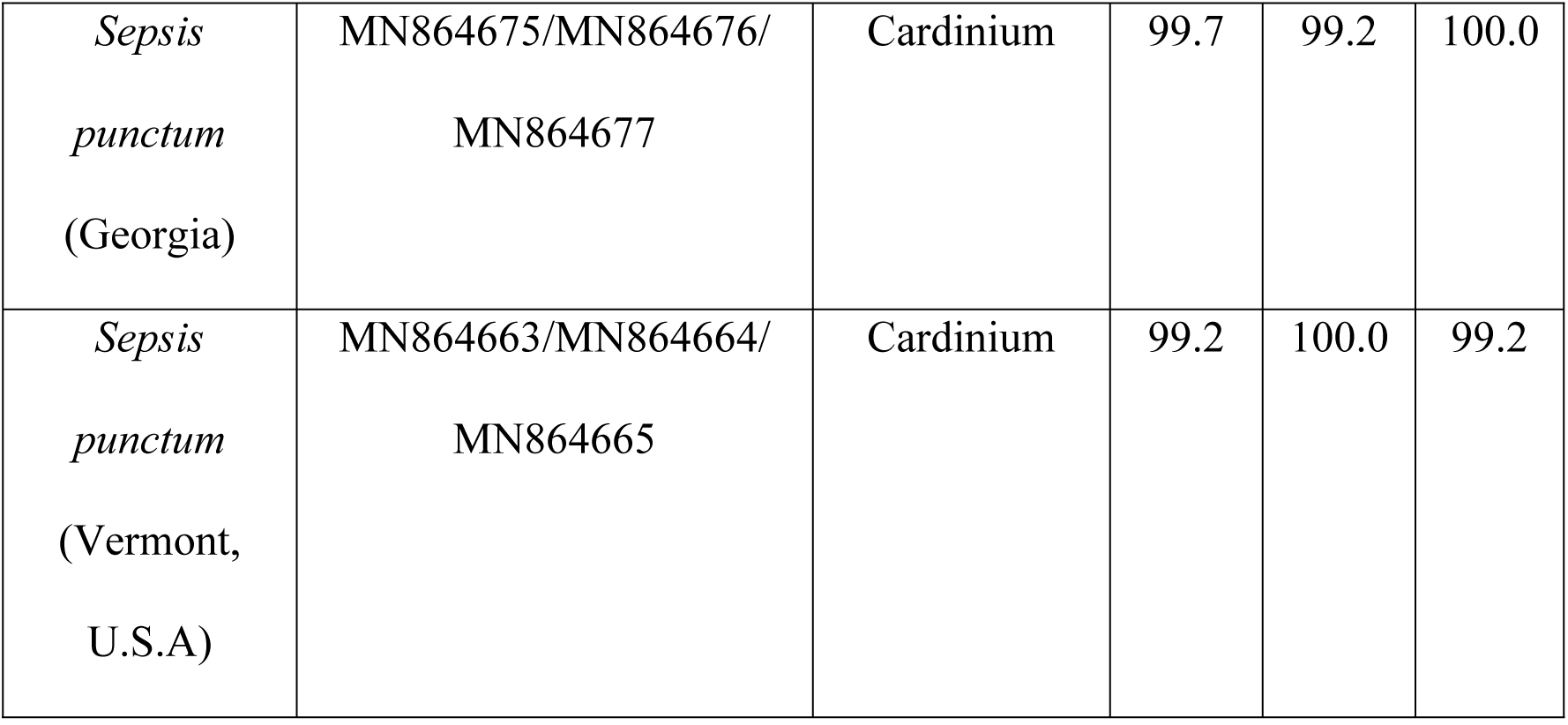
Pairwise sequence alignment of bacterial sequences obtained from the leg, egg or ovaries, and gut of the same fly, respectively. The Smith-Waterman algorithm was used to infer if the individual harboured the same strain of bacterium in all three tissues (e.g. Leg ↔Egg refers the alignment of amplicon sequence between the two specified tissues). The values represent percentage identity (%) between the respective DNA sequences.

## Discussion

There are very few published studies that investigate tissue-specific endosymbiont infections in arthropods (exceptions include (48,71,72)). Past studies focused on detecting tissue tropism and did not provide an optimized protocol to screen large volume of samples swiftly. Using the protocol, we conducted tissue-specific intra- as interspecific screening for two different fly families that are important model organisms in sexual selection research.

Additionally, there is no published tissue-specific screening protocol that simultaneously tested for co-infections and tissue tropism (10,11,41–44). In this study, we developed and optimised such a protocol. By using three sets of bacteria primers for each individual/ tissue, we recovered a high proportion of co-infections across both fly families (Table 1). Detection of such co-infections can alter the interpretations of studies looking at consequences of *Wolbachia* infections only. For instance, *Wolbachia* was known to interact with *Cardinium* to induce phenotypes different from single *Wolbachia*/ *Cardinium* infections (8, 73). Yet, single primer PCR was predominant in the past (43,44,74–76) and remains popular today (77–80). We suspect that the bias towards screening *Wolbachia* but not the other endosymbionts could be partly due to the inconvenience of conducting additional PCRs. By developing a multiplex PCR screening protocol, one or two less PCR reactions are required to screen for the three endosymbionts. Hopefully, this will encourage more researchers to screen for multiple endosymbionts.

There are, however, some limitations to our protocol. For instance, we found that some DNA extracted by QE could not be stored for a year and did not allow for successful amplification in subsequent PCRs. This is possibly due to the presence of active nucleases. If long-term storage is preferred over speed, we suggest using another DNA extraction method (such as phenol-chloroform) followed by duplex PCR. Next, the inability to dissect small specimens might hinder the feasibility of executing such a procedure. Also, contamination by the gut tissue might occur by accident. Then, researchers who are specimen-limited (e.g. working on rarer species) are likely only limited to whole insect PCR. Additionally, some insect species might harbour bacteria in the haemolymph and fat bodies (3,39,46) which are difficult to excise out during the dissection. However, the length of time needed to dissect and the likelihood of breaking the gastrointestinal tract decreases as the researcher becomes more experienced with practice. Lastly, our protocol could not discern between infected tissues if the bacteria were distributed throughout the host or if the bacterial gene was inserted into the host genome (47, 71). Previous research on tissue tropism detected infections in both somatic (including the gut) and germline tissues (48,71,72). In such cases, our protocol would only yield data on co-infection status.

An alternative to tissue-specific dissection is the fluorescence-based screening of *Wolbachia* using nucleic acid stain Syto-11 (81). While this method may be effective by allowing direct visualization of the endosymbiont in tissues, it is often more labour intensive, time-consuming and remains elusive to laboratories not equipped with confocal microscopy. Owing to the accessibility and convenience of PCR analysis, we recommend the tissue-specific protocol as an extension to current PCR methods for practical reasons.

We noticed that our triplex protocol was underpowered compared to the duplex despite the former producing repeatable results on positive controls. For multiplexes to evenly amplify all target DNA fragments, the balance between primer concentration and the amount of DNA is critical (58). Our optimization of a triplex PCR on the positive control worked because the primer concentrations were attuned specifically for the bacterial load in the sample. However, our data shows that the triplex is prone to false negatives, presumably because screened samples had a different volume of each bacteria compared to the positive control. The result was uneven amplification whereby signal for one or more bacteria may be too diluted to be observed on the gel. Such false negatives is a technical limitation of PCR multiplexing, (82, 83). However, Next-generation-sequencing using shotgun sequencing overcomes this limitation. In a previous study, such a metagenomic approach allowed for the detection of two *Wolbachia* strains (*wRi* and *wPip*) and *Spiroplasma* at low proportions in degraded *Amara alpina* samples (84). Nonetheless, a drawback of all NGS systems is the difficulty in optimizing sequencing depth (85). Specifically, a higher depth is required to detect bacterial DNA, which is lower in abundance than the host DNA. Yet, increased sequencing depth may produce more artefactual sequences (82).

### Single infection across tissues

A previous study documented *D. melanogaster* co-infected with *Wolbachia* and *Spiroplasma* harboured each bacteria in different tissues (7). This finding led to suspicions that different strains of the same bacterial species could be compartmentalized in different tissues. To investigate, we did a pairwise alignment between the bacterial 16S amplicons of the leg, egg and gut tissue belonging to the same individual. A total of six individuals, three *Drosophila* spp. and two *S. punctum* was examined. The results (Table 2) shows a high percentage of sequence identity (at least 98.8%, which is above the 97.5% criteria to be classified as the same strain (41)) between all aligned 355 bp bacterial 16S sequences. Also, the sequences did not show strong double peaks in the chromatogram, hence the presence of another low-density bacterial strain in the same sample was unlikely. This allows us to infer that at least some *Drosophila* spp. and sepsids harboured the same strain of *Wolbachia* and *Cardinium*, respectively.

### The implication of endosymbiont infections in drosophilid and sepsid flies

Despite being an increasingly popular model system for sexual size dimorphism and courtship behaviour studies, the black scavenger fly has yet to be screened for endosymbionts (23–25). This is the first study that has screened sepsid flies. *Wolbachia* infects about two-thirds of insect species with either 90% or 10% of individuals in each species being infected (86). Yet, we did not detect any *Wolbachia* in sepsids (Table 1). However, using primers for the *wsp* gene, we received positive bands of the expected size from sepsids of Singapore. Interestingly, only 5% of each sequence matched *wsp* in BLAST (Table S8). We hypothesize that these sepsids harbour an uncharacterized endosymbiont in which future taxonomic analysis may confirm.

The sepsids were largely infected with a less well-studied endosymbiont: *Cardinium* (Table 1 & S8). Every population, except for Tenerife, Wolverton, Tampines Eco-Green park and Pasir Ris, and 66.7 % of all individuals of sepsids harboured *Cardinium*. This high infection rate is surprising because a previous study screening 130 species in Dolichopodidae and Empidoidea found that only 10 species were infected with *Cardinium* (18).

Notably, horizontal transmission in *Rickettsia* was known to be transmitted sexually via the sperm storage organs (spermathecae) of females and the testes of males (72). It would be interesting to investigate if the *Rickettsia* in the sampled populations, which we have shown to be preferentially located in the gastrointestinal tract, were transmitted via the faecal-oral route. Also, a horizontal transfer may occur via ectoparasites such as mites (87) instead of the faecal-oral route. This might be feasible in sepsids which act as vehicles for mites from various dung pats (88). In our laboratory cultures, we commonly find mites on our sepsids. Thus, it may be prudent to incorporate the screening of the mites and other reproductive tissues besides ovaries or eggs in future studies.

As far as we know, this is the first screen of endosymbionts in wild populations of Singapore and the results suggest that a high infection rate in the drosophilids (30 individuals: *Wolbachia* - 86.7 %; *Cardinium* – 46.7 %; *Rickettsia* - 3.3 %). The BLAST results indicate that most of the local *Wolbachia* infections were identified to strain *wRi* which occurs naturally in *Drosophila simulans* (89). It was notable that the laboratory stock cultures exhibited a different rate of infection than the wild (42 individuals: *Wolbachia* – 23.1 %; *Cardinium* – 82.1 %; *Rickettsia* – 8.9 %). All but one sequenced *wsp* from the laboratory cultures were closely related to *Wolbachia* strain *w*Mel from *Drosophila melanogaster* (Table S9).

We observed multiple infections of the same individual by different endosymbionts (Table 1, S6-S7). Interestingly, *Rickettsia* was found only in samples with *Cardinium* (n=10, half of the infection were found in sepsids, the other half in drosophilids). In our screening of sepsids and *Drosophila* spp. (wild and stock), we discovered superinfections (infection by more than two endosymbionts) which occur naturally (73). One wild-caught *Drosophila* individual in Woodlands was infected by *Wolbachia*, *Cardinium*, *Rickettsia* (Table S7). Multiple infections must be considered when examining bacterial-induced phenotypes because the interaction between endosymbionts may result in additive or reductive effects on the intensity of traits such as cytoplasmic incompatibility, reduced bacterial load, and even introduce new traits such as feminization (7–9).

Aside from endosymbiont infections, it was notable that of the 25 out of 32 *Drosophila* spp. caught in Singapore were identified as *D. ananassae* (Table S1). This suggests that *D. ananassae* can be commonly found in Singapore and hence are likely candidates as wild-caught individuals for local research. Used in genetics and speciation studies (90–92), prospects of *D. ananassae* as models for behavioural studies exist as different populations vary in the oviposition behaviour (93). Although the possibility that the Singapore population of *D. ananassae* may not be indigenous, as they are cosmopolitan (93), it is still interesting to see if they differ from other populations in terms of genetics, evolution and behaviour.

### The paucity of studies on *Cardinium* infections

*Cardinium* in sepsids of 8 population were closely related to *Cardinium* in the whitefly *Bemisia tabaci* according to BLAST (Table S8). However, the BLAST results could have reflected the general lack of sequences of *Cardinium* in Genbank instead of the strong homology between the endosymbiont in sepsid and whitefly. We attempt to support this view through a search of all *Cardinium* sequences in Genbank that were derived from screening studies: only 756 sequences were found. This is a small number compared to the well-studied *Wolbachia* with a putative 204,546 sequences in GenBank. Therefore, there may be *Cardinium* more closely related to those in sepsids than that of *Bemisia tabaci* but remain undiscovered as *Cardinium* screenings were neglected. Also, among the 756 *Cardinium* sequences, only 317 was generated in conjunction with *Wolbachia* within the same study. This shows that most studies were interested in *Wolbachia* but not *Cardinium* despite studies showing their interactions can produce novel host phenotypes (94, 95).

## Conclusion

We believe that this protocol is time-efficient and cost-effective for researchers interested in studying the tissue-specific bacteria-induced phenotypes in insects. Interpreting the results of sexual selection studies can be confounded by the presence of bacteria such as *Wolbachia* (11). Using this protocol, future studies should be able to quickly screen their model species for bacterial infections before conducting any mating experiments.

## Acknowledgements

We would like to express gratitude to the staff of ReproLab, Hua Qian Hui, Sean Yap for their kind guidance. We are also thankful to Gwynne Lim, Denise Harjoko, Tiffany Lum, Tan Jia Wei, Pamela Kuan, and Toh Yi Peng, who collected *Drosophila* samples from their residences. Lastly, we owe our thanks to Jayanthi Puniamoorthy for assisting us in NGS sequencing and Dr John Ascher, Chui Shao Xiong and Toh Kai Xin for lending us their photography equipment. We thank Jack Werren (University of Rochester, NY) and Murad Ghamin (Volcanic enter, Israel) for sending us their positive samples. This study was partly funded by a start-up grant from the National University of Singapore (R154000A56133) as well as a Ministry of Education of Singapore Tier 1 grant (R154000A75114) awarded to the N. Puniamoorthy.

## Data availability statement

Raw datasets and chromatograms are deposited in Dryad (https://datadryad.org/stash/share/YSBMnX_Zrx16ZEoEnNO63gMEPTgbcQkjP20hNtkMgQY). Sequences are uploaded in Genbank as accessions MN864660-MN864706 and MN900912-MN900950.

## References

1. Hurst GDD, Crystal L. Reproductive Parasitism: Maternally lnherited Symbionts in a Biparental World. Cold Spring Harb Perspect Biol. 2015;7(a017699).

2. Werren J, Baldo L, Clark ME. *Wolbachia*: Master manipulators of invertebrate biology. Nat Rev Microbiol. 2008;6(10):741–51.

3. Goodacre SL, Martin OY. Modification of insect and arachnid behaviours by vertically transmitted endosymbionts: Infections as drivers of behavioural change and evolutionary novelty. Insects. 2012;3(1):246–61.

4. Werren J, Hurst G, Zhang W, Breeuwer J, Stouthamer R, Majerus M. Rickettsial Relative Associated with Male Killing in the Ladybird Beetle (*Adalia bipunctata*). J Bacteriol. 1994;176(2):388–94.

5. Hagimori T, Abe Y, Date S, Miura K. The First Finding of a *Rickettsia* Bacterium Associated with Parthenogenesis Induction Among Insects. Curr Microbiol. 2006;52(2):97–101.

6. Cordaux R, Bouchon D, Grève P. The impact of endosymbionts on the evolution of host sex-determination mechanisms. Trends Genet. 2011;27(8):332–41.

7. Goto S, Anbutsu H, Fukatsu T. Asymmetrical interactions between *Wolbachia* and *Spiroplasma* endosymbionts coexisting in the same insect host. Appl Environ Microbiol. 2006;72(7):4805–10.

8. Ros VID, Breeuwer JAJ. The effects of, and interactions between, Cardinium and Wolbachia in the doubly infected spider mite Bryobia sarothamni. Heredity (Edinb). 2009;102(4):413–22.

9. Curry MM, Paliulis L V., Welch KD, Harwood JD, White JA. Multiple endosymbiont infections and reproductive manipulations in a linyphiid spider population. Heredity (Edinb). 2015;115(2):146–52.

10. Martin OY, Puniamoorthy N, Gubler A, Wimmer C, Germann C, Bernasconi M V. Infections with the microbe *Cardinium* in the Dolichopodidae and other Empidoidea. J Insect Sci. 2013;13(1):47.

11. Martin OY, Gubler A, Wimmer C, Germann C, Bernasconi M V. Infections with *Wolbachia* and *Spiroplasma* in the Scathophagidae and other Muscoidea. Infect Genet Evol. 2012;12(2):315–23.

12. Moreau J, Bertin A, Caubet Y, Rigaud T. Sexual selection in an isopod with *Wolbachia*-induced sex reversal: Males prefer real females. Vol. 14, Journal of Evolutionary Biology. 2001. p. 388–94.

13. Vala F, Egas M, Breeuwer JAJ, Sabelis MW. *Wolbachia* affects oviposition and mating behaviour of its spider mite host. J Evol Biol. 2004;17(3):692–700.

14. De Crespigny FE, Pitt TD, Wedell N. Increased male mating rate in *Drosophila* is associated with *Wolbachia* infection. J Evol Biol. 2006;19(6):1964–72.

15. Arbuthnott D, Levin TC, Promislow DE. The impacts of *Wolbachia* and the microbiome on mate choice in *Drosophila melanogaster*. J Evol Biol. 2016;29(2):461–8.

16. Richard F-J. Symbiotic Bacteria Influence the Odor and Mating Preference of Their Hosts. Front Ecol Evol. 2017;5(143).

17. Hoffmann AA, Hercus M, Dagher H. Population dynamics of the *Wolbachia* infection causing cytoplasmic incompatibility in *Drosophila melanogaster*. Genetics. 1998;148(1):221–31.

18. Martin OY, Puniamoorthy N, Gubler A, Wimmer C, Bernasconi M V. Infections with *Wolbachia*, *Spiroplasma*, and *Rickettsia* in the Dolichopodidae and other Empidoidea. Infect Genet Evol. 2013;13:317–30.

19. Pont AC, Meier R. The Sepsidae (Diptera) of Europe. Vol. 37, Fauna Entomologica Scandinavica. Brill; 2002.

20. Puniamoorthy N, Su KFY, Meier R. Bending for love: Losses and gains of sexual dimorphisms are strictly correlated with changes in the mounting position of sepsid flies (Sepsidae: Diptera). BMC Evol Biol. 2008;8(155).

21. Puniamoorthy N, Ismail MRB, Tan DSH, Meier R. From kissing to belly stridulation: comparative analysis reveals surprising diversity, rapid evolution, and much homoplasy in the mating behaviour of 27 species of sepsid flies (Diptera: Sepsidae). J Evol Biol. 2009;22(11):2146–56.

22. Puniamoorthy N, Kotrba M, Meier R. Unlocking the “Black box”: internal female genitalia in Sepsidae (Diptera) evolve fast and are species-specific. BMC Evol Biol. 2010;10:275.

23. Dmitriew C, Blanckenhorn WU. The Role of Sexual Selection and Conflict in Mediating Among-Population Variation in Mating Strategies and Sexually Dimorphic Traits in *Sepsis punctum*. PLoS One. 2012;7(12):1–9.

24. Puniamoorthy N, Schäfer MA, Blanckenhorn WU. Sexual selection accounts for the geographic reversal of sexual size dimorphism in the dung fly, Sepsis punctum (Diptera: Sepsidae). Evolution (N Y). 2012;66(7):2117–26.

25. Rohner PT, Teder T, Esperk T, Lüpold S, Blanckenhorn WU. The evolution of male-biased sexual size dimorphism is associated with increased body size plasticity in males. Funct Ecol. 2017;32(2):581–91.

26. Lee HG, Kim YC, Dunning JS, Han KA. Recurring ethanol exposure induces disinhibited courtship in *Drosophila*. PLoS One. 2008;3(1).

27. Everaerts C, Farine JP, Cobb M, Ferveur JF. *Drosophila* cuticular hydrocarbons revisited: Mating status alters cuticular profiles. PLoS One. 2010;5(3):1–12.

28. Lin WS, Yeh SR, Fan SZ, Chen LY, Yen JH, Fu TF, et al. Insulin signaling in female *Drosophila* links diet and sexual attractiveness. FASEB J. 2018;32(7):3870–7.

29. Brown WD, Bjork A, Schneider K, Pitnick S. No evidence that polyandry benefits females in *Drosophila melanogaster*. Evolution (N Y). 2004;58(6):1242–50.

30. Bretman A, Fricke C, Chapman T. Plastic responses of male *Drosophila melanogaster* to the level of sperm competition increase male reproductive fitness. Proc R Soc B Biol Sci. 2009;276(1662):1705–11.

31. Gowaty P, Kim Y-K, Rawlings J, Anderson WW. Polyandry increases offspring viability and mother productivity but does not decrease mother survival in *Drosophila pseudoobscura*. Proc Natl Acad Sci. 2010;107(31):13771–6.

32. Chapman T, Liddle L, Kalb J, Wolfner M, Patridge L. Cost of mating in *Drosophila melanogaster* females is mediated by male accessory gland products. Nature. 1995;373(6511):241–4.

33. Miller G, Pitnick S. Sperm-Female Coevolution in *Drosophila*. Science (80-). 2002;298(5596):1230–3.

34. Gardiner A, Barker D, Butlin RK, Jordan WC, Ritchie MG. *Drosophila* chemoreceptor gene evolution: selection, specialization and genome size. Mol Ecol. 2008;17(7):1648–57.

35. O’Neill SL, Karr TL. Bidirectional incompatibility between conspecific populations of *Drosophila simulans*. Nature. 1990;348(6297):178–80.

36. Vavre F, Fleury F, Lepetit D, Fouillet P, Boulétreau M. Phylogenetic Evidence for Horizontal Transmission of *Wolbachia* in Host-Parasitoid Associations. Mol Biol Evol. 1999;16(12):1711–23.

37. Rasgon JL, Gamston CE, Ren X. Survival of *Wolbachia* pipientis in cell-free medium. Appl Environ Microbiol. 2006;72(11):6934–7.

38. Caspi-Fluger A, Inbar M, Mozes-Daube N, Katzir N, Portnoy V, Belausov E, et al. Horizontal transmission of the insect symbiont *Rickettsia* is plant-mediated. Proc R Soc B Biol Sci. 2012;279(1734):1791–6.

39. Frost CL, Pollock SW, Smith JE, Hughes WOH. *Wolbachia* in the Flesh: Symbiont Intensities in Germ-Line and Somatic Tissues Challenge the Conventional View of *Wolbachia* Transmission Routes. PLoS One. 2014;9(7).

40. Li S-J, Ahmed MZ, Lv N, Shi P-Q, Wang X-M, Huang J-L, et al. Plant-mediated horizontal transmission of *Wolbachia* between whiteflies. ISME J. 2017 Apr 9;11(4):1019–28.

41. Zhou W, Rousset F, O’neill S. Phylogeny and PCR based classification of Wolbachia strains using wsp gene sequences. Proc R Soc L B Biol Sci. 1998;265(December 1997):509–15.

42. Jeyaprakash A, Hoy MA. Long PCR improves *Wolbachia* DNA amplification: *wsp* sequences found in 76% of sixty-three arthropod species. Insect Mol Biol. 2000;9(4):393–405.

43. Pourali P, Roayaei Ardakani M, Jolodar A, Razi Jalali MH. PCR screening of the Wolbachia in some arthropods and nematodes in Khuzestan province. Iran J Vet Res. 2009;10(3):216–22.

44. Simões PM, Mialdea G, Reiss D, Sagot MF, Charlat S. *Wolbachia* detection: an assessment of standard PCR protocols. Mol Ecol Resour. 2011;11(3):567–72.

45. Baldo L, Ayoub NA, Hayashi CY, Russell JA, Stahlhut JK, Werren JH. Insight into the routes of *Wolbachia* invasion: High levels of horizontal transfer in the spider genus *Agelenopsis* revealed by *Wolbachia* strain and mitochondrial DNA diversity. Mol Ecol. 2008;17(2):557–69.

46. Pietri JE, DeBruhl H, Sullivan W. The rich somatic life of *Wolbachia*. Microbiologyopen. 2016;5(6):923–36.

47. Kondo N, Nikoh N, Ijichi N, Shimada M, Fukatsu T. Genome fragment of *Wolbachia* endosymbiont transferred to X chromosome of host insect. Proc Natl Acad Sci U S A. 2002;99(22):14280–5.

48. Doudoumis V, Tsiamis G, Wamwiri F, Brelsfoard C, Alam U, Aksoy E, et al. Detection and characterization of *Wolbachia* infections in laboratory and natural populations of different species of tsetse flies (genus *Glossina*). BMC Microbiol. 2012;12(12(Suppl 1):S3):1–13.

49. Dunning Hotopp JC, Clark ME, Oliveira DCSG, Foster JM, Fischer P, Muñoz Torres MC, et al. Widespread lateral gene transfer from intracellular bacteria to multicellular eukaryotes. Science (80-). 2007;317(5845):1753–6.

50. Semiatizki A, Weiss B, Bagim S, Rohkin-Shalom S, Kaltenpoth M, Chiel E. Effects, interactions, and localization of *Rickettsia* and *Wolbachia* in the house fly parasitoid, Spalangia endius. Microb Ecol. 2020;

51. Mateos M, Castrezana SJ, Nankivell BJ, Estes AM, Markow TA, Moran NA. Heritable endosymbionts of *Drosophila*. Genetics. 2006;174(1):363–76.

52. Kaur R, Martinez J, Rota-Stabelli O, Jiggins FM, Miller WJ. Age, tissue, genotype and virus infection regulate *Wolbachia* levels in *Drosophila*. Mol Ecol. 2020;29(11):2063–79.

53. Rohner PT, Blanckenhorn WU, Puniamoorthy N. Sexual selection on male size drives the evolution of male-biased sexual size dimorphism via the prolongation of male development. Evolution (N Y). 2016;70(6):1189–99.

54. Markow TA, O’Grady PM. Drosophila A Guide to Species Identification and Use. Elsevier; 2006. 145–153 p.

55. Lee Kong Chian Natural History Museum. The Biodiversity of Singapore [Internet]. [cited 2018 Oct 7]. Available from: https://singapore.biodiversity.online/

56. Ratnasingham S, Hebert PDN. BARCODING, BOLD : The Barcode of Life Data System (www.barcodinglife.org). NPR Mol Ecol Notes. 2007;7:355–64.

57. Brown S, Chen Y-W, Wang M, Clipson A, Ochoa E, Du M-Q. PrimerPooler: automated primer pooling to prepare library for targeted sequencing. Biol Methods Protoc. 2017;2(1):1–10.

58. Sint D, Raso L, Traugott M. Advances in multiplex PCR: Balancing primer efficiencies and improving detection success. Methods Ecol Evol. 2012;3(5):898–905.

59. Meier R, Wong W, Srivathsan A, Foo M. $1 DNA barcodes for reconstructing complex phenomes and finding rare species in specimen-rich samples. Cladistics. 2015;32(1):100–10.

60. Technelysium. ChromasPro. Technelysium; 2018.

61. Weeks AR, Velten R, Stouthamer R. Incidence of a new sex-ratio-distorting endosymbiotic bacterium among arthropods. Proc R Soc B Biol Sci. 2003;270(1526):1857–65.

62. NCBI Resource Coordinators. Database resources of the National Center for Biotechnology Information. Nucleic Acids Res. 2016;44:D7–19.

63. Sánchez R, Serra F, Tárraga J, Medina I, Carbonell J, Pulido L, et al. Phylemon 2.0: A suite of web-tools for molecular evolution, phylogenetics, phylogenomics and hypotheses testing. Nucleic Acids Res. 2011;39(SUPPL. 2):470–4.

64. Edgar RC. MUSCLE: Multiple sequence alignment with high accuracy and high throughput. Nucleic Acids Res. 2004;32(5):1792–7.

65. Capella-Gutiérrez S, Silla-Martínez JM, Gabaldón T. trimAl: A tool for automated alignment trimming in large-scale phylogenetic analyses. Bioinformatics. 2009;25(15):1972–3.

66. Kumar S, Stecher G, Li M, Knyaz C, Tamura K. MEGA X: Molecular Evolutionary Genetics Analysis across Computing Platforms. Mol Biol Evol. 2018;35(6):1547–9.

67. RStudio Team. RStudio: Integrated Development for R. [Internet]. Boston: RStudio, PBC; 2020. Available from: https://www.rstudio.com/

68. Sweet L. nonpar: A Collection of Nonparametric Hypothesis Tests [Internet]. 2017 [cited 2019 Aug 20]. Available from: https://cran.r-project.org/web/packages/nonpar/index.html

69. Mangiafico S. rcompanion: Functions to Support Extension Education Program Evaluation [Internet]. 2019. Available from: https://cran.r-project.org/web/packages/rcompanion/index.html

70. Stackebrandt, Erko; Ebers J. Taxonomic parameter revised: tarnished gold standards. Microbiol Today. 2006;8(4):6–9.

71. Cheng Q, Ruel TD, Zhou W, Moloo SK, Majiwa P, O’Neill SL, et al. Tissue distribution and prevalence of *Wolbachia* infections in tsetse flies, Glossina spp. Med Vet Entomol. 2000;14(1):44–50.

72. Brumin M, Levy M, Ghanim M. Transovarial transmission of *Rickettsia* spp. and organ-specific infection of the whitefly *Bemisia tabaci*. Appl Environ Microbiol. 2012;78(16):5565–74.

73. Zhao DX, Chen DS, Ge C, Gotoh T, Hong XY. Multiple Infections with *Cardinium* and Two Strains of *Wolbachia* in The Spider Mite *Tetranychus phaselus* Ehara: Revealing New Forces Driving the Spread of *Wolbachia*. PLoS One. 2013;8(1):1–9.

74. Zchori-Fein E, Perlman S. Distribution of the bacterial symbiont *Cardinium* in arthropods. Mol Ecol. 2004;13(7):2009–16.

75. Enigl M, Schausberger P. Incidence of the endosymbionts *Wolbachia*, *Cardinium* and *Spiroplasma* in phytoseiid mites and associated prey. Exp Appl Acarol. 2007;42(2):75–85.

76. Duron O, Bouchon D, Boutin S, Bellamy L, Zhou L, Engelstädter J, et al. The diversity of reproductive parasites among arthropods: *Wolbachia* do not walk alone. BMC Biol. 2008;6(27):1–12.

77. Duplouy A, Brattström O. *Wolbachia* in the Genus Bicyclus: a Forgotten Player. Microb Ecol. 2018;75(1):255–63.

78. Leggewie M, Krumkamp R, Badusche M, Heitmann A, Jansen S, Schmidt-Chanasit J, et al. *Culex torrentium* mosquitoes from Germany are negative for *Wolbachia*. Med Vet Entomol. 2018;32(1):115–20.

79. Goindin D, Cannet A, Delannay C, Ramdini C, Gustave J, Atyame C, et al. Screening of natural *Wolbachia* infection in *Aedes aegypti*, *Aedes taeniorhynchus* and *Culex quinquefasciatus* from Guadeloupe (French West Indies). Acta Trop. 2018;185:314–7.

80. Kulkarni A, Yu W, Jiang J, Sanchez C, Karna AK, Martinez KJL, et al. *Wolbachia* pipientis occurs in *Aedes aegypti* populations in New Mexico and Florida, USA. Ecol Evol. 2019;9(10):6148–56.

81. Casper-Lindley C, Kimura S, Saxton DS, Essaw Y, Simpson I, Tan V, et al. Rapid fluorescence-based screening for *Wolbachia* endosymbionts in *Drosophila* germ line and somatic tissues. Appl Environ Microbiol. 2011;77(14):4788–94.

82. Ekblom R, Galindo J. Applications of next generation sequencing in molecular ecology of non-model organisms. Heredity (Edinb). 2011;107(1):1–15.

83. Rennstam Rubbmark O, Sint D, Horngacher N, Traugott M. A broadly applicable COI primer pair and an efficient single-tube amplicon library preparation protocol for metabarcoding. Ecol Evol. 2018;8(24):12335–50.

84. Heintzman PD, Elias SA, Moore K, Paszkiewicz K, Barnes I. Characterizing DNA preservation in degraded specimens of *Amara alpina* (Carabidae: Coleoptera). Mol Ecol Resour. 2014;14(3):606–15.

85. Rennstam Rubbmark O, Sint D, Cupic S, Traugott M. When to use next generation sequencing or diagnostic PCR in diet analyses. Mol Ecol Resour. 2019;19(2):388–99.

86. Hilgenboecker K, Hammerstein P, Schlattmann P, Telschow A, Werren JH. How many species are infected with *Wolbachia*? – statistical analysis of current data. FEMS Microbiol Lett. 2008 Apr;281(2):215–20.

87. Jaenike J, Polak M, Fiskin A, Helou M, Minhas M. Interspecific transmission of endosymbiotic *Spiroplasma* by mites. Biol Lett. 2007;3(1):23–5.

88. Kiontke K. The Phoretic Association of Diplogaster coprophila Sudhaus and Rehfeld, 1990 (Diplogastridae) From Cow Dung With Its Carriers, in Particular Flies of the family sepsidae. Vol. 42, Nematologica. Brill; 1996. 354–366 p.

89. Turelli M, Cooper BS, Richardson KM, Ginsberg PS, Peckenpaugh B, Antelope CX, et al. Rapid Global Spread of wRi-like *Wolbachia* across Multiple *Drosophila*. Curr Biol. 2018;28(6):963–71.

90. Coyne JA, Orr HA. “Patterns of Speciation in *Drosophila*” Revisited. Evolution (N Y). 1997;51(1):295–303.

91. Fay J, Chung-I W. Hitchhiking Under Positive Darwinian Selection. Genetics. 2000;155(3):1405–13.

92. Drosophila 12 Genomes Consortium. Evolution of genes and genomes on the Drosophila phylogeny. Nature. 2007;450(7167):203–18.

93. Singh BN. Population and behaviour genetics of *Drosophila ananassae*. Genetica. 1996;97(3):321–9.

94. White JA, Kelly SE, Cockburn SN, Perlman SJ, Hunter MS. Endosymbiont costs and benefits in a parasitoid infected with both *Wolbachia* and *Cardinium*. Heredity (Edinb). 2011;106(4):585–91.

95. Zhu LY, Zhang KJ, Zhang YK, Ge C, Gotoh T, Hong XY. *Wolbachia* strengthens *Cardinium*-induced cytoplasmic incompatibility in the spider mite *Tetranychus piercei* McGregor. Curr Microbiol. 2012;65(5):516–23.

